# Determining the factors driving selective effects of new nonsynonymous mutations

**DOI:** 10.1101/071209

**Authors:** Christian D. Huber, Bernard Kim, Clare D. Marsden, Kirk E. Lohmueller

**Author notes:** **To whom correspondence should be addressed:** Christian D. Huber, Department of Ecology and Evolutionary Biology, University of California, Los Angeles, 621 Charles E. Young Drive South, Los Angeles, CA 90095-1606, Kirk E. Lohmueller, Department of Ecology and Evolutionary Biology, University of California, Los Angeles, 621 Charles E. Young Drive South, Los Angeles, CA 90095-1606.

## Abstract

The distribution of fitness effects (DFE) of new mutations is a fundamental parameter in evolutionary genetics^1–3^. While theoretical models have emphasized the importance of distinct biological factors, such as protein folding^4^, back mutations^5^, species complexity^6,7^, and mutational robustness^8^ at determining the DFE, it remains unclear which of these models can describe the DFE in natural populations. Here, we show that the theoretical models make distinct predictions about how the DFE will differ between species. We further show that humans have a higher proportion of strongly deleterious mutations than *Drosophila melanogaster*. Comparing four categories of theoretical models, only Fisher’s Geometrical Model (FGM) is consistent with our data. FGM assumes that multiple phenotypes are under stabilizing selection, with the number of phenotypes defining a complexity of the organism. It suggests that long-term population size and cost of complexity drive the evolution of the DFE, with many implications for evolutionary and medical genomics.

## Main text

The distribution of fitness effects (DFE) is a fundamental paraymeter in evolutionary genetics because it quantifies the amount of deleterious, neutral, and adaptive genetic variation in a population^3^. Despite the importance and considerable study of the DFE^1–3^, the biological factors determining the DFE in different species remain elusive. Several theoretical models propose different mechanisms for the evolution of the DFE^4–6,8,9^. While each of these models has a reasonable theoretical basis as well as some support from experimental evolution studies or microbial studies, which model best explains differences in the DFE between species has not yet been determined. Nor have these models been tested with genetic variation data from natural populations in higher organisms. Although experimental evolution studies in laboratory organisms might more closely match the assumptions of the models being tested, natural populations may provide different qualitative results due to increased resolution to measure weakly deleterious mutations and unnatural selection pressure in the laboratory^1,10^.

Importantly, the five theoretical models for the evolution of the DFE predict that the DFE will differ between species with different levels of organismal complexity and long-term population size (Fig. 1). Here we leverage this prediction to test which theoretical model best explains the evolution of the DFE by comparing the DFE in natural population of humans and Drosophila. To do this, we utilized polymorphism data of a sample of 112 individuals from Yoruba in Ibadan, Nigeria (YRI) from the 1000 Genomes project^11^ and 197 African *Drosophila melanogaster* lines from the *Drosophila* Population Genomics Project^12^. We summarize the polymorphism data by the folded site frequency spectrum (SFS), which represents the number of variants at different minor allele frequencies in the sample (Supplementary Fig. 1A). Because population history can also affect patterns of polymorphism, we first use the synonymous SFS to estimate demographic models separately in each species. We infer that the population size of YRI and Drosophila expanded 2.3-fold 5,500 generations ago and 2.7-fold 500,000 generations ago, respectively (Supplementary Table 1). Note that demographic estimates from synonymous sites are biased by selection on linked sites^13^, but that this bias does not affect performance of the DFE estimation^14^ (see Methods).

**Figure 1.**
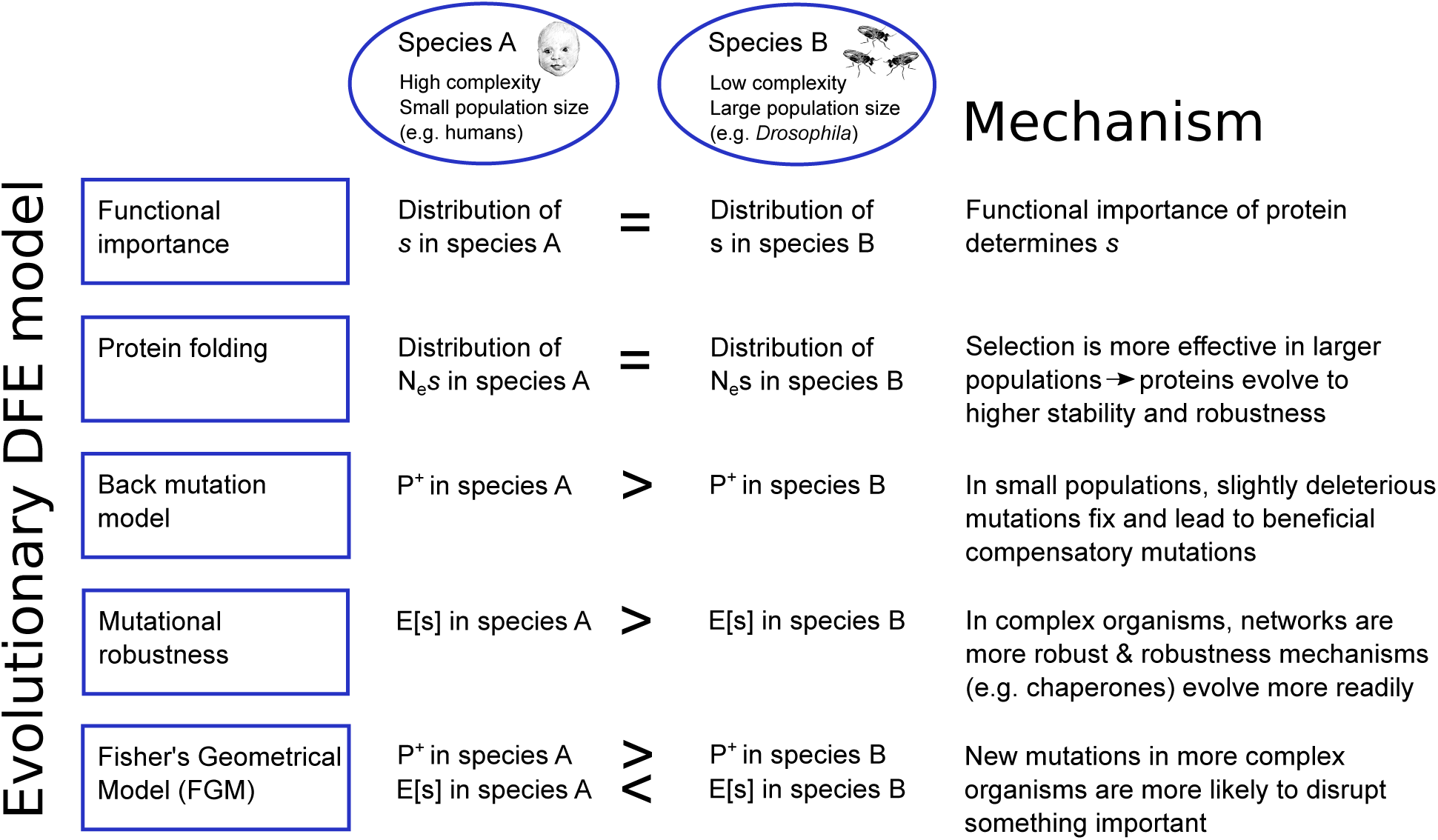
Overview of main predictions of five theoretical models regarding DFE differences between two species. Here, *P*^+^ is the proportion of slightly beneficial mutations, E[*s*] is the average selection coefficient, and *N*_e_ is the effective population size. Note that more negative (i.e. lower) E[*s*] implies more strongly deleterious mutations. Subscript A refers to species A, subscript B refers to species B. See Supplementary Note 2 for more details.

Conditional on the estimated demographic parameters, we estimate the DFE for new nonsynonymous mutations in both species using the nonsynonymous SFS. In short, our approach utilizes the fact that more deleterious mutations segregate in lower numbers and at lower frequencies than less deleterious or neutral mutations. Thus, we do not directly quantify the deleteriousness of any specific mutation, but indirectly summarize the fitness effects over many sites by estimating the parameters of a DFE that fits the SFS. It was shown that as long as the demographic parameters estimated from the synonymous data can fit the synonymous SFS, then the inference of the DFE for the nonsynonymous sites remains unbiased, even when the true data include background selection, population growth, and non-modeled population structure^14–16^. Here, we compare the estimates of the DFE from the two species in a novel likelihood ratio test framework that accounts for differences in recent demographic history between the two species (see Methods). Briefly, we assume that the DFE follows a gamma distribution, and find that a model where each species has its own shape and scale parameters fits the SFSs for the two species significantly better than a model where the parameters are constrained to be the same across both species (Likelihood Ratio Test (LRT) statistic Λ=920; df=2, *P*<10^−16^). This result holds even when making different assumptions about the mutation rate, selection on synonymous sites, as well as when omitting singleton variants (Supplementary Note 1; Supplementary Table 2; Supplementary Table 3). Examination of the maximum likelihood gamma distribution shows that *Drosophila* has a much higher proportion of weakly deleterious and nearly neutral mutations with selection coefficient *s* (a measure of the relative fitness effect of a mutation) > −10^−4^ than do humans (Fig. 2D). The proportion of strongly deleterious mutations with *s* < −10^−3^ is significantly larger in humans (55%) than in *Drosophila* (5%). Thus, our results provide statistical support for humans and *Drosophila* having different DFEs (of *s*) that cannot be explained by differences in population size or demography between the species. To evaluate the robustness of our finding to the assumed functional form of the DFE, we tested a range of different distributions other than the gamma or log-normal, as well as a nonparametric discretized distribution. We consistently find that mutations are on average more deleterious in humans than in *Drosophila* (Supplementary Note 1, Supplementary Fig. 8, Supplementary Fig. 14, and Supplementary Table 4).

**Figure 2.**
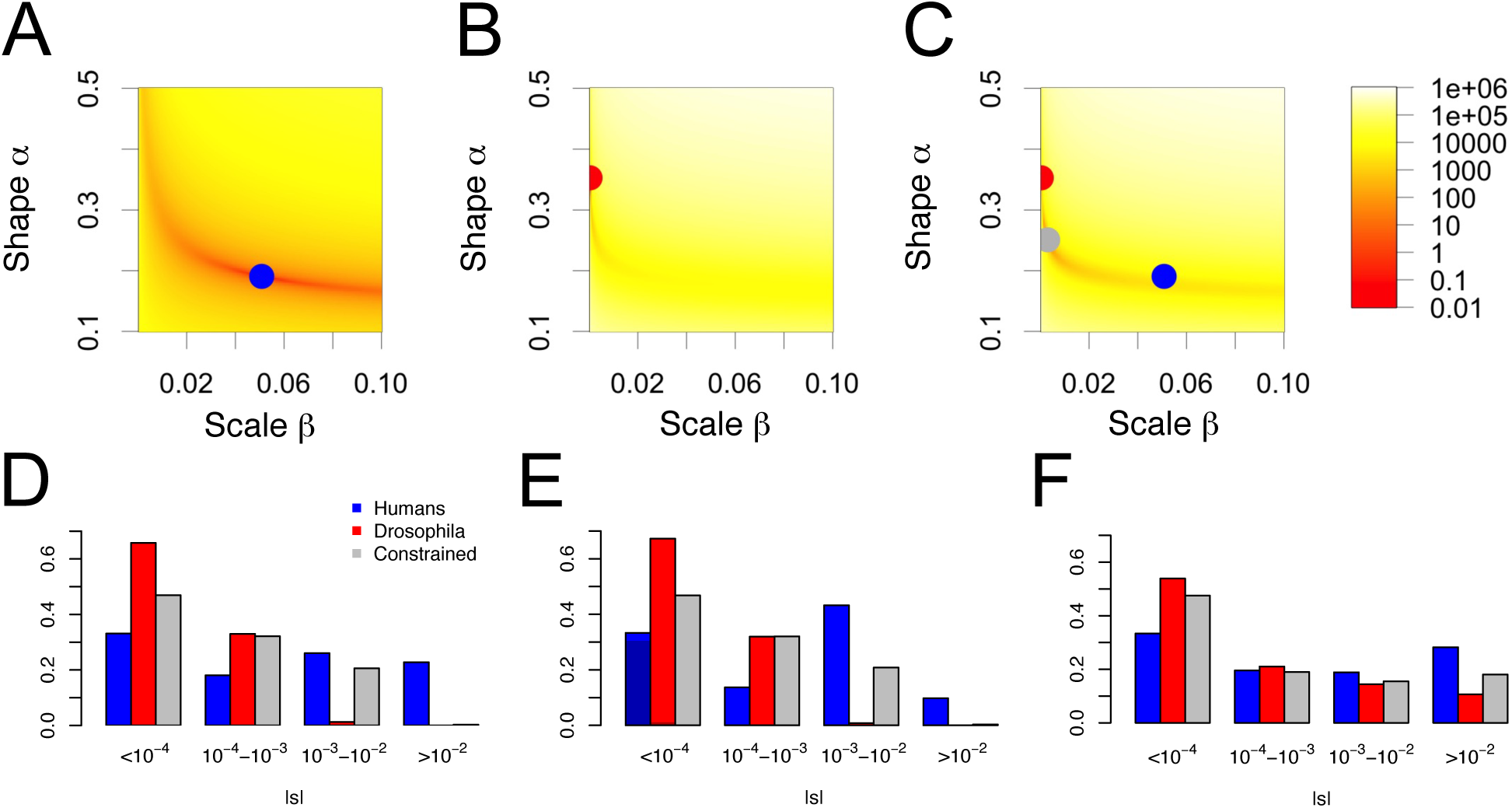
Testing the null hypothesis of the same distribution of *s* in both species. The log-likelihood surface for the shape and scale parameters of a gamma-distributed DFE(*s*) for (A) humans, (B) *Drosophila*, and (C) both datasets combined (constrained model). Colors from yellow to red indicate the difference in log-likelihood of that set of parameter values compared to the MLE (see color scale). E.g. orange indicates parameters ∼ 100 log-likelihood units below the MLE. Proportions of mutations for various ranges of |*s*| are computed from the estimated (D) gamma distribution, (E) mixture of gamma distribution with neutral point mass, and (F) log-normal distribution. The grey bars indicate the proportions under the null hypothesis of the same distribution of *s* in both species (constrained model). Darker colors in (E) reflect the estimated proportions of neutral mutations.

Because a variety of demographic, statistical, and numerical biases can confound LRTs using the SFS, we evaluated the performance of our statistical approach by analyzing simulated datasets. Specifically, we performed forward-in-time simulations that include realistic levels of linkage disequilibrium and background selection (Supplementary Note 1). When we estimated the DFE from the simulations of the full model, the estimates were unbiased (Fig. 3A,B). This suggests that the size change model fit to synonymous polymorphisms successfully controls for the effects of background selection (Supplementary Fig. 3; see also ref.^13^). As expected, the null distribution of Λ derived from simulations under the constrained model is broader than the chi-square distribution with two degrees of freedom (Fig. 3C). However, all of the 300 Λ values that we simulated were smaller than 34, suggesting the probability of seeing a Λ value bigger than 920 is substantially less than 0.33% under the null. Since selective sweeps were suggested to be a major determinant of genetic diversity in *Drosophila*^17^, we also examined the effect of recurrent selective sweeps on our inference. In line with other studies^14^, we found that selective sweeps do not significantly bias our DFE estimates when correcting for the effect of demography using the observed SFS at neutral sites (Supplementary Fig. 9 and Supplementary Note 1). In summary, a combination of confounding factors cannot account for our findings of different DFEs between human and *Drosophila* (see also Supplementary Note 1).

**Figure 3.**
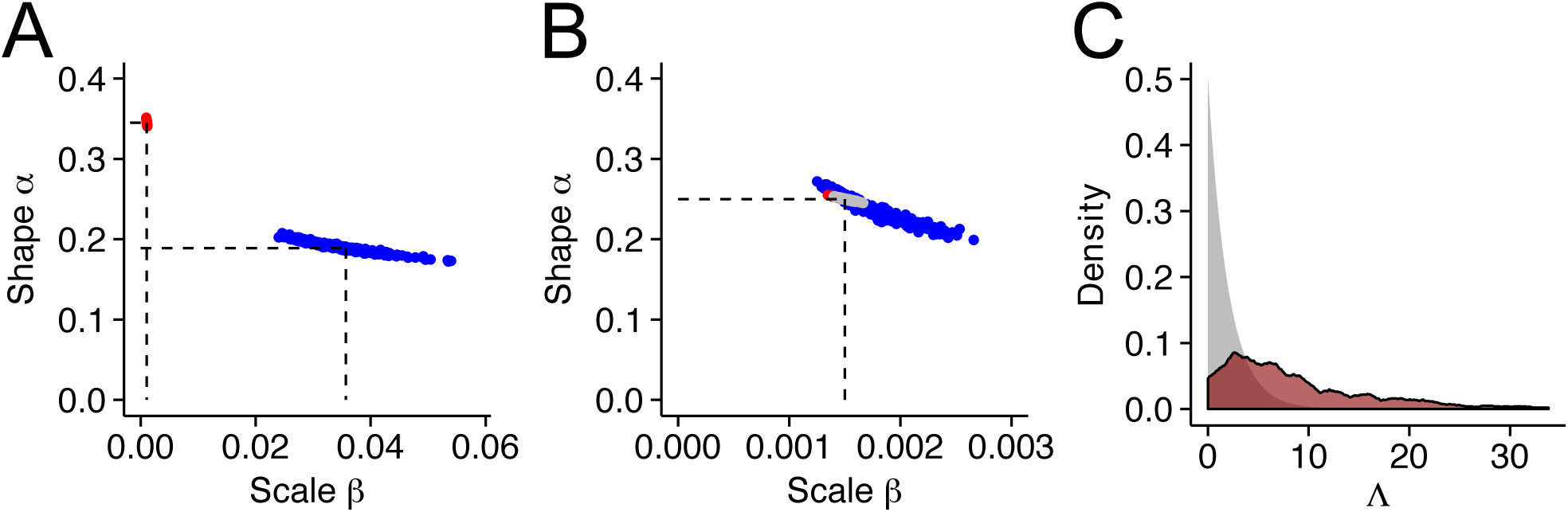
Estimates of the shape and scale parameters of a gamma DFE from 300 simulations of human (blue) and *Drosophila* (red) data. (A) Estimates from simulations under the alternative hypothesis (H1), i.e. assuming maximum likelihood parameters in both species (dashed lines). Results show that we can retrieve the right parameters. (B) Estimates from simulations under the null hypothesis (H0), i.e. assuming a single set of parameters in both species (dashed lines). In grey are the estimation results using data from both species simultaneously, assuming H0 is correct. Results show that, under H0, we correctly retrieve the same set of parameters for both species. (C) The expected (grey) and simulated (dark red) null distribution of the test statistic Λ = −2*log(L_Constrained,max_/L_Full,max_) for testing the null hypothesis of no difference in shape and scale parameters between humans and *Drosophila*.

Next we tested whether differences in the DFE between species vary across functional categories of genes. First, when restricting our analysis to a strict ortholog set, the significant difference in the DFE between humans and *Drosophila* remained (Λ = 7,369, p < 10^−16^). Further, the parameter estimates were very similar between the two sets of genes (Supplementary Fig. 5A, Supplementary Table 2, Supplementary Table 3). To examine the effect of gene expression on the DFE, we classify genes into sets with different gene expression profiles (Supplementary Fig. 1B; Supplementary Fig. 2; see Methods). Overall, we found that, though the shape parameter varies between tissue specific and broadly expressed genes, the average selection coefficient E[*s*] is about 50–80 fold more negative for humans than for *Drosophila*, regardless of the overall expression level or tissue specificity of the genes (Supplementary Fig. 12A). These results suggest that although the DFE may vary across genes with distinct expression profiles, differences in expression alone are insufficient to account for the observed differences in the DFE between the two species.

Having established that common confounders and differences in gene expression cannot account for the differences in the DFE between species, we next examined which of the four theoretical models can explain the differences. The first model, the protein stability model, predicts that much of the selection pressure involves maintaining the thermodynamic stability of proteins. This model predicts that the distribution of *N_e_s* is gamma distributed^18^ and independent of the effective population size (*N_e_*) when at equilibrium^4^ (see Supplementary Note 2 for specific assumptions). Thus, this model predicts that *N_e_s* is the same across taxa. However, in contrast to this prediction, we found that a model with different *N_e_s* distributions in each species fit the data significantly better than a model where *N_e_s* was constrained to be the same in both species (Λ = 22,000, p < 10^−16^; Supplementary Fig. 4; Supplementary Fig. 6), consistent with previous results^16^. Comparing this LRT statistic to the null distribution obtained from forward simulations similar to those discussed above suggests that such a large LRT statistic is highly incompatible with a model that assumes the same gamma (or lognormal) *N_e_s* distribution in both species (p < 0.0033). Thus, our data do not support protein stability models as the driving force in the evolution of the DFE between species.

The second model, the back-mutation model, predicts that there is a category of weakly advantageous mutations that restore fitness after deleterious mutations become fixed^19^. The back-mutation model predicts that in small populations, the proportion of slightly beneficial mutations is greater than in large populations, because more slightly deleterious mutations can become fixed in small populations, leading to more opportunities for new beneficial back mutations (Supplementary Note 2). Using this logic, Piganeau and Eyre-Walker^20^ (see also Rice et al.^5^) derived a formula for the equilibrium DFE as a function of population size. When we estimate the parameters in the model from our data in our framework, we found an unrealistically large effective population size in *Drosophila* (5.2x10^19^). Further, we inferred distinct parameters of the effect size distribution (the distribution of |*s*|) in the two species (Supplementary Table 4) such that the average effect size E[|*s*|] of a mutation in humans is about 80 fold larger than in *Drosophila*, which is inconsistent with the predictions of the back-mutation model (see also Supplementary Fig. 8B). Although the Piganeau and Eyre-Walker model fits well within both species, it falls short in providing an evolutionary or mechanistic explanation for a large difference in E[|*s*|] between species.

The third model, the mutational robustness model, postulates that more robust, or complex, organisms have, on average, less deleterious mutations^6,8^. Here, more complex organisms have a greater ability to compensate and buffer the effects of deleterious mutations (Supplementary Note 2). Note that complexity can be hard to define and quantify in a biologically and evolutionarily meaningful way. However, a number of biological factors suggest humans are more complex than *Drosophila*. Such factors include a larger number of genes, a larger number of proteins and protein-protein interactions^21^, and likely also a larger number of cell types^22^ in humans than in *Drosophila*. Mutational robustness model predict greater mutational robustness in humans than in *Drosophila* because of the higher complexity and the smaller effective population size of humans compared to *Drosophila*. However, inconsistent with this prediction, we have shown that humans have a 50-80 fold more negative value of E[*s*] than *Drosophila*, and a larger proportion of strongly deleterious mutations with *s* < −0.001 (Fig. 2D-F). Further, robustness models predict that less pleiotropic mutations are more deleterious, since the smaller effective complexity of such mutations impedes the evolution of robustness^23^. Assuming that broadly expressed genes are more pleotropic than tissue-specific genes, we observe that tissue-specific genes have less negative estimates of E[*s*] than broadly expressed genes (Supplementary Fig. 12A). In other words, more pleiotropic mutations tend to be more deleterious. This finding is inconsistent with predictions from the robustness models. However, while our results suggest that mutational robustness mechanisms are not the main driver of differences in the DFE across species, this finding is not necessarily at odds with previous work on these models. The clearest empirical evidence for an increase of mutational robustness by selection comes from experimental evolution studies of viruses and bacteria^24,25^. Viruses and bacteria have large mutation rates and population sizes. The specific mechanism that promotes robustness in such organisms may not be applicable to higher organisms with smaller population mutation rates^26^. Our results suggest that if mutational robustness mechanisms play a role in shaping the DFE of higher organisms, they do not compensate for other factors that increase the deleteriousness of mutations in humans compared to *Drosophila*.

The fourth model, Fisher’s Geometric Model (FGM) represents phenotypes as points in a multidimensional phenotype space and fitness is a decreasing function of the distance to the optimal phenotype^6^. The dimensionality of the phenotype space is termed “complexity”. FGM makes three predictions that we test with our data (Supplementary Note 2). The first prediction is that more complex organisms, like humans, have more deleterious mutations than *Drosophila*, since mutations are more likely to disrupt something important in a complex organism than in a simple one^27^ (see Supplementary Note 3 for assumptions that go into this prediction). Indeed, this prediction is well supported by our data because the average selection coefficient E[*s*] is estimated to be 50-80 times more negative in humans than in *Drosophila*. To further validate this finding in a larger phylogenetic context, we analyzed polymorphism data from mouse (*Mus musculus castaneus*) and yeast (*Saccharomyces paradoxus*). Although sample size is one order of magnitude smaller, we replicate the pattern of increasing deleteriousness of mutations with increasing complexity (Fig. 4A, Supplementary Table 5). Second, smaller populations are predicted to have a larger proportion of beneficial mutations due to increased fixation of deleterious mutations in smaller populations when populations are in equilibrium (drift load^28^). Note that population size here refers to long-term effective population size, thus it could be affected by background selection and selective sweeps as well as demographic processes. To test this prediction, we estimated the parameters for the DFE based on FGM. Formulas have been derived for the DFE assuming the population is at an arbitrary distance from the optimal phenotype (eq. 8 of Lourenço et al.^28^ and eq. 5 in Martin and Lenormand^7^), or assuming mutation-selection-drift equilibrium (eq. 15 of Lourenço et al.^28^). We found that the equilibrium DFE fits just as well or better than the non-equilibrium versions (Supplementary Table 4), suggesting that in both populations, most genes are close to equilibrium and that the DFE is a function of *N_e,long-term_*. Further, in humans, the equilibrium Lourenço DFE shows a significantly better fit over the plain gamma DFE (Supplementary Table 4), with a *N_e,iong-term_* of 2100 (95% CI: 1653 - 2546). Note that this value of *N_e,long-term_* is of the same order of magnitude as the ancestral population size estimated from synonymous sites (6,600). This is surprising since the estimate of *N_e,long-term_* is not based on neutral diversity, but on the degree of maladaptation due to drift load that results in some proportion of beneficial compensatory mutations in the DFE. Thus, it is estimated from the predicted effect of drift load on the nonsynonymous SFS and likely reflects a much larger time-span than the estimate from the synonymous SFS. In *Drosophila*, fitting the equilibrium Lourenço model led to a similar fit as the plain gamma DFE (Supplementary Table 4). Further, the large *N_e,long-term_* (8.4×10^7^) estimated here is also similar to that estimated from the neutral synonymous sites (2.8×10^6^). The fact that long-term population sizes inferred under FGM are consistent with previous estimates from genetic variation data suggests that this prediction of FGM is satisfied by our data. Third, FGM predicts that more pleotropic mutations will show smaller variation in *s*. As before, we use gene expression breath as a proxy for pleiotropy. We found that the shape parameter (α) of the gamma distribution is smaller for tissue-specific genes than for broadly expressed genes (Fig. 4B). The shape parameter is inversely related to the coefficient of variation (CV) of the selection coefficient: CV(*s*)=1/sqrt(α). Thus, the smaller shape parameter indicates a larger CV(*s*) and is consistent with the idea that mutations in tissue-specific genes are less pleiotropic than in broadly expressed genes. Similar conclusions were derived by explicitly estimating pleiotropy from fitting the Lourenço DFE to the data (Supplementary Fig. 13).

**Figure 4.**
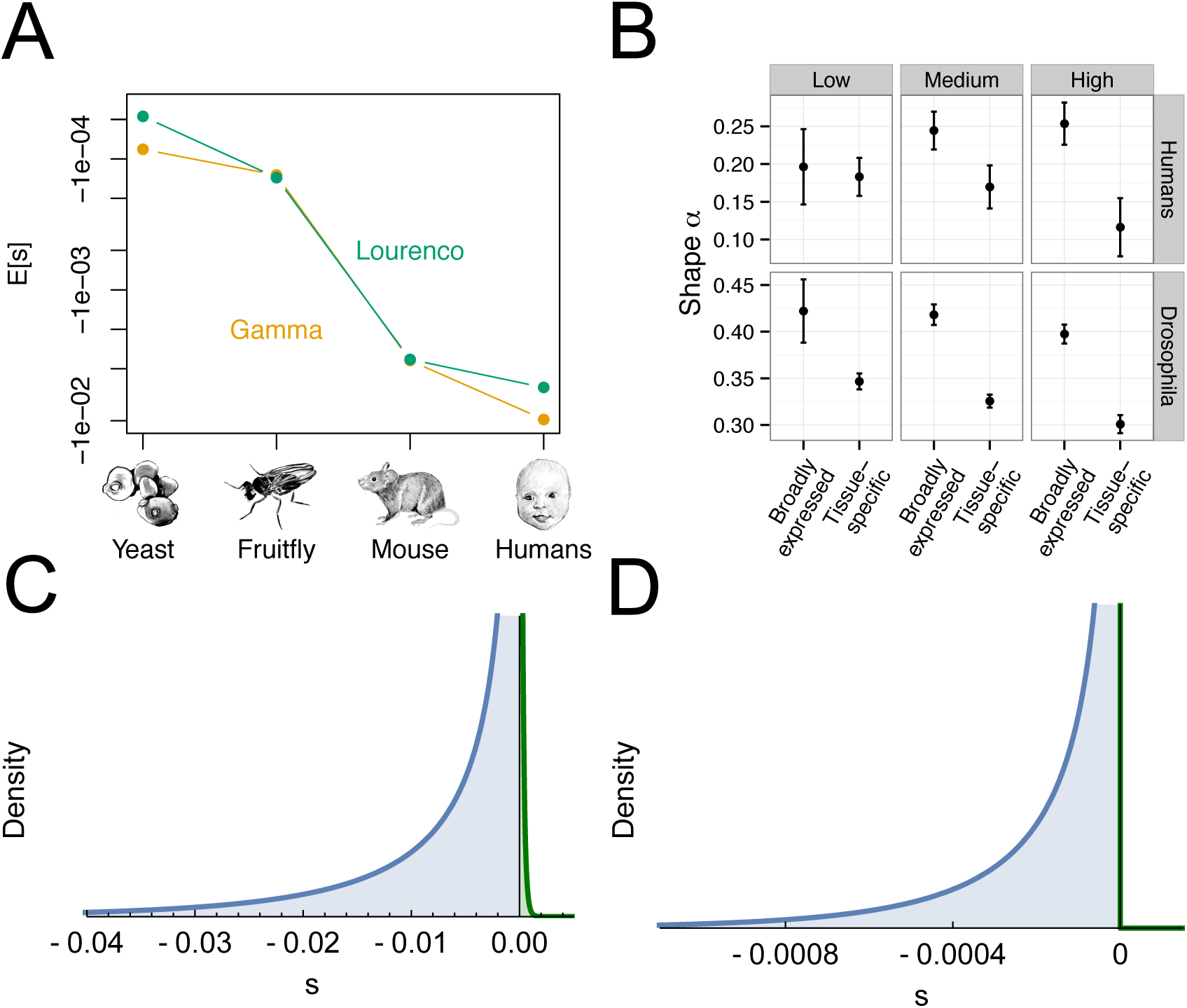
Empirical support for FGM. (A) Both under the gamma DFE and the Lourenço et al.^28^ DFE, estimated average deleteriousness of mutations increases as a function of organismal complexity. (B) The shape parameter of the gamma DFE depends on the breadth of gene expression. Tissue-specific genes have a smaller shape parameter (α) than broadly expressed genes, indicating less pleiotropy in tissue-specific genes. (C,D) By fitting the DFE of Lourenço et al. we can model slightly beneficial mutations in the DFE (green) that are thought to compensate for fixed deleterious mutations in small population size species. We find support for a larger proportion of slightly beneficial mutations in the DFE of (A) humans than in (B) *Drosophila*.

In sum, all three predictions made by FGM are supported by our data. We conclude that FGM is a viable model to explain differences in the DFE between species and genes. Under this model, species complexity as well as distance of the population to the fitness optimum, modulated by long-term population size, are the key drivers of the DFE of new amino-acid mutations. Note that many essential elements of protein evolution are captured by FGM^29^, where many molecular phenotypes (not just protein stability) are under stabilizing selection^30^. Thus, although we reject a simple protein stability model determining the DFE, this should not be taken to mean that general principles of protein evolution do not play a role in determining the DFE.

Our findings have implications for important aspects of evolutionary genetics. First, FGM allows us to estimate the proportion of new mutations that are adaptive. When assuming FGM, we estimate that 15% of new nonsynonymous mutations in humans are beneficial. The majority (96%) of these beneficial mutations have small selection coefficients, with *s* < 0.0005 (Fig. 4C). In *Drosophila*, however, the model including positive selection had a similar fit as the plain gamma DFE (Supplementary Table 4), and only 1.5% of new mutations are beneficial (Fig. 4D). This finding appears to be at odds with previous studies of adaptive evolution in these two species. The proportion of amino acid substitutions that fixed due to positive selection was estimated to be larger in *Drosophila* (50%) than in humans (10–20%), using a McDonald-Kreitman (MK) approach^2,31^. More generally, our results suggest that inferences of the amount of adaptive evolution considering fixed substitutions may be fundamentally and qualitatively different from those considering new mutations. Additionally, the amount of positive selection in the human genome has been recently debated^32,33^. After controlling for background selection, Enard et al.^32^ found that, in humans, estimates of the amount of adaptive evolution from MK approaches may be severe underestimates. Their results instead argue that there may be many small-scale adaptive steps in humans, i.e. many weak selective sweeps that are only detectable when averaging across many instances. Such a mode of adaptation is in fact predicted by FGM for organisms with high complexity^34^, but see ref.^35^.

Second, a varying DFE over phylogenetic timescales has implications for understanding the overdispersed molecular clock^36^. The substitution rate of deleterious mutations relative to the rate of neutral evolution is a function of the compound parameter *N_e_s*^37^. Thus, not only phylogenetic changes in *N*_*e*_ but also changes in *s* may contribute to overdispersion. Our results suggest that changes in the distribution of *s* are coupled with changes in population size and complexity. For example, the larger complexity of humans is supposed to reduce the nonsynonymous divergence along the human lineage to lower values than what would be expected from the two orders of magnitude population size difference to *Drosophila*. Accurate characterization of the DFE from many species across the tree of life will enable a direct test of the contribution of changing DFEs to the dispersion of the molecular clock.

Lastly, our results have implications for assessing the biological function of sequences using evolutionary information. The comparative genomics paradigm postulates that biologically important regions of the genome are constrained across long evolutionary times^38^. This implies that *s* for a particular sequence is determined by the biological importance of the sequence and that *s* remains constant over time. If, as our work suggests, selection coefficients change over time as a consequence of species complexity and long-term population size, this could result in important sequences not showing the prototypical signatures of conservation, leading to such sequences being missed by comparative approaches. Further, it suggests that complexity and population size are important factors to consider when deciding which species to utilize in future comparative genomic studies.

## Acknowledgments

We thank Bridgett vonHoldt, Tanya Phung and Sebastian Matuszewski for comments that greatly improved the manuscript, Toni I. Gossmann for providing access to site frequency spectrum data from *Mus musculus castaneus* and *Saccharomyces paradoxus*, and Maria T. Huber for the drawings in Fig. 1 and 4. C.D.H., C.D.M, and K.E.L. were supported by a Searle Scholars Fellowship and an Alfred P. Sloan Research Fellowship in Computational & Molecular Biology to K.E.L.

## Author Contributions

Conceptualization, C.D.H. and K.E.L.; Methodology, C.D.H. and K.E.L., Software, C.D.H. and B.K.; Validation, C.D.H. and B.K.; Formal Analysis, C.D.H.; Resources, K.E.L.; Data Curation, C.D.M. and B.K.; Writing – Original Draft, C.D.H. and K.E.L.; Writing – Review & Editing, C.D.H., B.K., C.D.M., and K.E.L.; Visualization, C.D.H.; Supervision, K.E.L.

## Online Methods

### Data

We used published next generation sequencing data sets to extract the synonymous and nonsynonymous SFS. For humans, we used the sample of 112 individuals from Yoruba in Ibadan, Nigeria (YRI) from the 1000 Genomes Project^11^. We downloaded the 1000 Genomes phase 3 dataset from the 1000 Genomes ftp site (ftp://ftp.1000genomes.ebi.ac.uk/vol1/ftp/phase3/, accessed Sept 2014). Using information in the sample information PED file, related individuals were removed and for each trio or family group only the mother and father were used. The SNPs were also filtered for whether they were within the exome capture array region and in the strict mask part of the human genome, as defined by the 1000 Genomes Project. The genotypes of YRI individuals were extracted and annotated using the SeattleSeq annotation pipeline (http://snp.gs.washington.edu/SeattleSeqAnnotation138/). For *Drosophila melanogaster*, we used the DPGP phase 3 data of a sample of 197 lines originating from Zambia, Africa^12^. We accessed whole genome genotype data for the 197 genomes from the Pool lab (http://johnpool.net/genomes.html). These data were provided in non-standard vcf format (vcf sites file, downloaded August 2014), therefore we first converted these to a standard vcf format with the BDGP5.75 genome as the reference using a custom python script. We then merged all the individual vcf files and removed any sites with evidence of identity by descent or admixture using the masking package provided by the Pool lab. Only the 2L, 2R, 3L and 3R chromosome arms were used in our analyses. We then conducted variant annotation using SnpEeff v3.6 using the BDGP5.75 database.

We filtered both datasets for sites with sample size > 99 and down-sampled all sites with larger sample size than 100 to a sample size of 100 using the hypergeometric probability distribution. Further, we selected only sites that were in exons and computed an exon length *L_exon,i_* for each gene *i*. The nonsynonymous and synonymous sequence length (*L_NS_*, *L_S_*) depends on the transition/transversion ratio and CpG mutational bias. We assumed a transition:transversion ratio of 2:1 in *Drosophila*^39,40^ and 3:1 for human exons^41,42^. Further, we assumed a 10x mutational bias at CpG sites in humans, but no such effect in *Drosophila*^43^. This leads to multipliers of *L_NS_* = 2.85 × *L_S_* in *Drosophila*, and *L_NS_* = 2.31 × *L_S_* in humans. We calculated the synonymous and nonsynonymous SFS, and the respective sequence lengths (*L_NS,i_*, *L_S,i_*), for each gene *i*. For all further inference, we used the folded SFS to avoid correcting for misidentification of the ancestral state. Ancestral misidentification could lead to unwanted and difficult to control biases^44^.

To study the effect of gene expression on the DFE, we used two recent gene expression datasets from humans^45^ and *Drosophila*^46^ that provide mRNA expression level estimates in 27 and 29 different tissues, respectively. For both datasets, we transformed the ‘fragments per kilobase of exon per million fragments’ (FPKMs) by computing log(FPKM+1) and quantile normalizing this value over all tissues using ‘normalize.quantiles’ of the R package ‘preprocessCore’, resulting in an expression level S. We computed τ as a measure of tissue specificity for each gene: 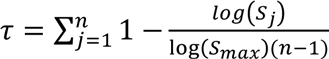. Here, n is the number of tissues, S_j_ is the expression level in tissue j and S_max_ is the largest expression level over all tissues. We used τ to classify genes as tissue specific (τ > 0.6) or broadly expressed (τ < 0.4). We further classified genes as low (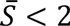), intermediate (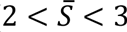) and highly expressed (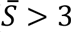), where 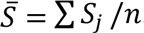. This classification leads to strongly different gene expression profiles between classes (Supplementary Fig. 2), but still enough data in every class to be able to reliably estimate the DFE (θ_s_ > 100 in humans and θ_s_ > 900 in *Drosophila*).

To infer the DFE in *Mus musculus castaneus* (mouse) and *Saccharomyces paradoxus* (yeast), we used the synonymous and nonsynonymous SFS data from Gossmann et al.^47^. Our estimates of proportions of mutations in different *N_e_s* bins (Supplementary Fig. 10) are concordant with what has been preported previously^47–49^. We then used mutation rate estimates for yeast^50^ and mouse^51^, respectively, to estimate *N_e_* and transform the DFE from *N_e_s* to *s*.

### Estimating demography and DFE

We used the software δaδi^52^ to infer the parameters of a single size change model using the synonymous site frequency spectrum (SFS) under the Poisson Random Field framework^53^. In this framework, the multinomial likelihood quantifies how well the empirical SFS fits to an expected SFS that is derived from specific demographic parameters^52^. Assume that Θ_D_ is a vector of demographic parameters (i.e., time and strength of a population size change), X_i_ is the count of SNPs with frequency i, P_i_ is the proportion of SNPs at frequency i, θ is the population mutation rate, and n is the sample size. The distribution of allele frequency q in the population (g[q|ΘD]) can be computed by numerically solving the diffusion approximation to the Wright-Fisher model, and can also incorporate selection^2,52,54^. We used δaδi^52^ to numerically approximate g[q|ΘD]. Further, the expected number of SNPs at frequency i in a sample of size n is 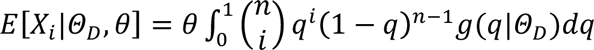. The relative proportion of SNPs at frequency i can then be calculated as 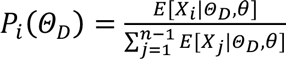, and the formula for the multinomial likelihood is 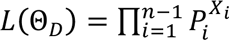. To derive the maximum likelihood estimate of Θ_D_ 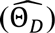 we maximized *L*(Θ_D_).

We used the Poisson likelihood instead of the multinomial likelihood to estimate the vector of parameters of the DFE (Θ_DFE_). We found that this strongly improves the precision of the scale parameter of the gamma distribution compared to using the multinomial likelihood since the Poisson likelihood uses information from both the absolute number of SNPs as well as the curvature of the SFS^2,15^. Note however that we do not make use of fixed differences to an outgroup. Including information from fixed differences hardly improves inferring the DFE of deleterious mutations^55^, which are the main focus of our paper. The likelihood of Θ_DFE_ was thus calculated as 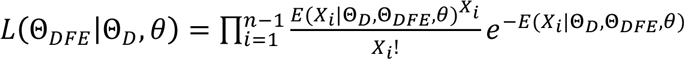. We set Θ_D_ here to the maximum likelihood estimates of the demographic parameters 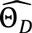, and *θ* to the nonsynonymous population mutation rate *θ_NS_* = 4*N_e_µL_NS_*. We estimated *θ_NS_* from *θ_S_* by accounting for the difference in synonymous and nonsynonymous sequence length.

The formula of the Poisson likelihood depends on *E*(*X_i_*|Θ_*D*_, Θ_*DFE*_, *θ*), i.e. on the expected SFS given the demography, *θ_NS_* and some distribution of *N_e_s* with parameters Θ_*DFE*_. However, δaδi only allows computing the expected SFS *E*(*X_i_*|Θ_*D*_, *N_e_s*, θ) for a single selection coefficient *N_e_s* (and some arbitrary demography). Thus, we extend δaδi’s functionality by computing the expected SFS for a grid of 1000 *N_e_s* values on an exponentially distributed grid between −15000 and −10^−4^. This set of site frequency spectra is further used to calculate the expected SFS for an arbitrary distribution of N_e_s values. This is done by numerically integrating over the respective spectra weighted by the gamma distribution. The numerical integration was done using the ‘numpy.trapz’ function as implemented in δaδi. Due to numerical instabilities for strongly skewed distributions, we did not integrate all the way towards 0, but computed the weight of *N_e_s* values between –10^−4^ and 0 and added the product of this weight with the neutral SFS to the expected SFS. Mutations with *N_e_s* < −15000 are expected not to contribute to the SFS since they are strongly selected against. Our approach allows us to estimate the parameters of any arbitrary distribution of *N_e_s* values. We implemented the gamma distribution, log-normal distribution, the formula of Piganeau and Eyre-Walker ^20^, eq. 7, assuming gamma distributed effect sizes, and the formula of Lourenço et al.^28^, eq. 15. The formula of Lourenço et al.^28^ provides an explicit solution to the DFE for Fisher’s geometrical model under fitness equilibrium. It is a function of three parameters: population size, effect size, and the average number of phenotypes affected by a mutation (pleiotropy). The DFE of Lourenço et al.^28^ and Piganeau and Eyre-Walker^20^ are distributions with some proportion of slightly beneficial mutations. In models with some proportion of beneficial mutations, those mutations are expected to segregate in the population and thus influence both the shape of the SFS and the absolute number of SNPs. We use this expectation to infer the full DFE (beneficial plus deleterious mutations) from the SFS, similar to Tataru et al.^14^. To do this, we also integrate over beneficial mutations with *N_e_s* from 0 to 15000. Numerical optimization is used to find the parameters of the DFE distribution that maximize the poisson likelihood. For this optimization step, we use the BFGS algorithm as implemented in the ‘optimize.fmin_bfgs’ function of scipy. To avoid finding local optima, we repeated every estimation approach (for both the simulations and the real data) from 50 uniformly distributed random starting parameters. Standard errors were based on the Hessian matrix of the log-likelihood function, numerically computed at the maximum likelihood estimates using the ‘Hessian.hessian’ function of δaδi^52^. They were computed as the square root of the diagonal elements of the inverse of the negative Hessian matrix^56^. Confidence intervals were approximated as plus/minus two times the standard errors, except where specified otherwise.

Note that population genetic methods for estimating the DFE from the SFS can only estimate the composite parameter of selection coefficient *s* with effective population size *N*_*e*_, since the effect of selection on the SFS depends on *N_e_s* and not *s* alone. However, the distribution of *s* can be derived from the distribution of *N_e_s* by scaling it by 1/*N_e_* (e.g. multiplying the scale parameter of a gamma distribution of *N_e_s* by 1/*N_e_*). Fitting the demographic model to the synonymous SFS provided an estimate of θ_S_ = 4*N_e_µL_S_* for synonymous sites, where µ is the neutral per base-pair mutation rate and *L_S_* is the synonymous sequence length. Using this formula, we estimated *N_e_* by setting the neutral mutation rate to either 2.5×10^−8^ for humans and 1.5×10^−9^ for *Drosophila* (phylogenetic estimates^57–59^) or to 1.5×10^−8^ for humans and 3×10^−9^ for *Drosophila* (current estimates ^58,60,61^). Note that when partitioning our data into different gene categories and estimating the DFE for each category separately, we also allow for a different ancestral *N_e_* and demography estimates in those categories to control for different levels of background selection in different genomic regions^62,63^.

### Statistical test for different DFEs between two species

We used the SFS from polymorphism data from two species, A (*X_i,A_*) and B (*X_j,B_*), to test whether the DSE differs between those two species. First, we estimated the demographic model parameters of both species (Θ*_D,A_*, Θ*_D,B_*) as outlined above. Second, we assumed that the DFE in both species follows a gamma distribution with the shape parameter *α* and scale parameter *β*. We used a Poisson composite likelihood function, where the SFS at nonsynonymous SNPs in species A is treated as being independent of that from species B, which is reasonable for distantly related species with little incomplete lineage sorting^55^. Then, the likelihood function for the parameters is:

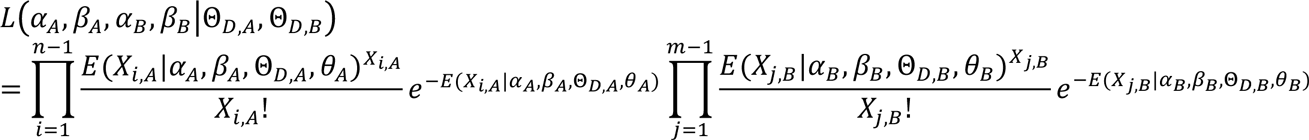

Here, *n* and *m* are the sample size of species A and species B, respectively. We will test whether the shape (*α*) and scale (*β*) parameters in species A differ from those in species B. To do this, we propose the following likelihood ratio test (LRT):

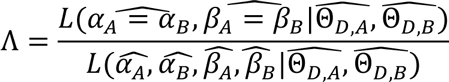

The null hypothesis (constrained model) is that *α_A_* = *α_B_* and *β_A_* = *β_B_*. The full model allows for *α_A_* ≠ *α_B_* and *β_A_* ≠ *β_B_*. We optimized the likelihood function under both the null and full models as outlined above. Importantly, in all cases, we conditioned on the demographic parameters in each population, thus accounting for differences in population history. Asymptotically, Λ follows a chi-square distribution with 2 degrees of freedom, due to the two additional free parameters in the full model compared to the constrained model. Simulations were used to test how well the usual asymptotic theory applies in this situation. The test is not limited to comparing the parameters of a gamma distribution of two species, but can be extended to any DFE distribution (e.g. log-normal), and any number of species, in a straightforward way. The degree of freedom of the chi-square null distribution is *p^*^k-p*, where *p* is the number of parameters of the distribution, and *k* is the number of species.

### Forward simulations

To compute the null distribution of the likelihood ratio test statistic, Λ, we performed forward simulations under the estimated demographic models for humans and *Drosophila*. Selection coefficients for nonsynonymous mutations were drawn from a gamma distribution with shape and scale parameters estimated from the constrained model (i.e., α_H_ = α_D_ and β_H_ = β_D_). We assume a spatial distribution of selected elements that reflects the empirical distribution of coding and conserved non-coding (CNC) sequence in the genome. Further, we simulate varying recombination across the genomes that is based on empirical high-resolution recombination maps^64,65^. Mutations in CNC regions are assumed to be selected with gamma distributed selection coefficients taken from Torgerson et al.^66^ for humans and Casillas et al.^67^ for *Drosophila*. The exon element ranges where taken from GENCODE v14^68^ for humans and BDGP 6.79 FlyBase gene annotation^69^ for *Drosophila*. To define CNC ranges in both species, we used predicted conserved elements by phastCons^70^, downloaded from the UCSC genome browser. All forward simulations were carried out using the simulation software ‘SLiM’^71^. For both species, we simulated under a single size change model with the empirically estimated parameters (Supplementary Table 1). Since *Drosophila* has a prohibitively large population size for forward simulations, we simulated both species with an ancestral effective population size of 10,000 and scaled mutation rate, recombination rate, selection coefficients and demographic parameters accordingly^72^. To assess power, we performed a different set of simulations assuming the gamma DFE parameter estimates from the full model (Supplementary Table 2).

Further, to allow quasi genome-wide simulations, we followed a bootstrapping approach by first simulating 1000 × 7 Mb large regions that were selected randomly from the respective genome. We then selected a centered 3 Mb window from the simulated 7 Mb region and discharged the rest of the sequence to remove edge effects, notably the lower strength of background selection at the edges^73^. From those 1000 × 3 Mb regions, we resampled until we arrive at a full genome data set, i.e. synonymous and nonsynonymous SFS that are similar in size to the actual data. That way, we simulated data of 300 independent genomes. In both species, the simulations resulted in considerable amounts of background selection, with average reduction in neutral diversity in the 7Mb region of 10% in humans and 12% in *Drosophila*. For each simulated genome data we first estimated the demographic model from the synonymous SFS and then the DFE parameters from the nonsynonymous SFS conditional on the estimated demographic parameters.

